# Towards generalizable prediction of antibody thermostability using machine learning on sequence and structure features

**DOI:** 10.1101/2022.06.03.494724

**Authors:** Ameya Harmalkar, Roshan Rao, Jonas Honer, Wibke Deisting, Jonas Anlahr, Anja Hoenig, Julia Czwikla, Eva Sienz-Widmann, Doris Rau, Austin Rice, Timothy P. Riley, Danqing Li, Hannah B. Catterall, Christine E. Tinberg, Jeffrey J. Gray, Kathy Y. Wei

**Author notes:** A.H and R.R share equal author contribution.

## Abstract

Over the last three decades, the appeal for monoclonal antibodies (mAbs) as therapeutics has been steadily increasing as evident with FDA’s recent landmark approval of the 100th mAb. Unlike mAbs that bind to single targets, multispecific biologics (bsAbs) with their single-chain variable fragment (scFv) modules have garnered particular interest owing to the advantage of engaging distinct targets. Despite their exquisite specificity and affinity, the relatively poor thermostability of these scFv modules often hampers their development as a potential therapeutic drug. In recent years, engineering antibody sequences to enhance their stability by mutations has gained considerable momentum. As experimental methods for antibody engineering are time-intensive, laborious, and expensive, computational methods serve as a fast and inexpensive alternative to conventional routes. In this work, we show two machine learning methods - one with pre-trained language models (PTLM) capturing functional effects of sequence variation, and second, a supervised convolutional neural network (CNN) trained with Rosetta energetic features - to better classify thermostable scFv variants from sequence. Both these models are trained over temperature-specific data (TS50 measurements) derived from multiple libraries of scFv sequences. In this work, we show that a sufficiently simple CNN model trained with energetic features generalizes better than a pre-trained language model on out-of-distribution (blind) sequences (average Spearman correlation coefficient of 0.4 as opposed to 0.15). Further, we demonstrate that for an independent mAb with available thermal melting temperatures for 20 experimentally characterized thermostable mutations, these models trained on TS50 data could identify 18 residue positions and 5 identical amino-acid mutations showing remarkable generalizability. Our results suggest that such models can be broadly applicable for improving the biological characteristics of antibodies. Further, transferring such models for alternative physico-chemical properties of scFvs can have potential applications in optimizing large-scale production and delivery of mAbs or bsAbs.

## 1. Introduction

Monoclonal antibodies (mAbs) represent a large class of therapeutic agents, with more than 100 FDA-approved products marketed in the US (www.antibodysociety.org). Despite their widespread prevalence in drug development, mAbs are limited in biological scope because they bind only a single target. Multispecific biologics (bsAbs) engaging more than one target or epitope on the same target are of growing importance for accessing novel, therapeutically relevant pathways and mechanisms of action. In recent years, several multispecific biologics are approved for use and many more are in clinical and preclinical development (1, 2). A common building block for the construction of multispecific biologics is the single-chain variable fragment (scFv), consisting of the target-engaging variable heavy chain (VH) linked to the variable light chain (VL) via a flexible linker. Multispecific format platforms such as the BiTE (3), IgG-scFv (4), and XmAb (5) incorporate scFv modules. Although scFvs are prevalent in multispecific biologic candidates, they may display sub-optimal physical properties relative to conventional mAbs and generally require sequence modifications to produce a developable asset. One property that is used to gauge the potential developability of a scFv module or scFv-containing multispecific is thermostability – scFv candidates are experimentally screened and/or optimized for thermostability to identify suitable modules (6, 7). However, these experiments are resource intensive and time consuming. Accurate computational methods to predict scFv thermostability from primary amino acid sequence for scFv candidate selection/deselection (and to predict mutations to guide thermostability engineering efforts) would be invaluable to multispecific drug development.

Over the past decade, the use of computational tools to predict stability-enhancing mutations has gained considerable momentum, albeit with limited success. Protein consensus design, a state-of-the-art approach, uses phylogenetic information from multiple sequence alignments (MSAs) to obtain the most frequent amino acid for a residue position. But, these residues have improved thermostability in only 50% of the cases, with even poorer performance for antibody sequences with highly conserved framework regions. Structure-based approaches, such as PROSS (8) and AbLIFT (9), have used thermodynamic energies (Rosetta ΔΔG) and structural information (CDR-dependent PSSMs) to improve binding and stability, however their application to scFvs is often limited by the relative scarcity of structural data. Alternatively, machine learning approaches have employed large datasets such as ProTherm (10), that collate mutant effects on protein stability from mesophilic and thermophilic sequences, to predict thermostability (11, 12). Unfortunately, the plethora of publicly available thermostability data excludes mAb or scFv sequences, limiting their generalization. Predicting thermally-enhancing scFv sequences or designing thermostable mutations within approved scFv candidates is still a challenge. Currently, there is a need for computational approaches to leverage sequence information to predict biophysical attributes, such as thermostability, with high accuracy and generalizability enabling efficient protein design. We address this critical need by equipping unsupervised and supervised learning approaches over a thermostability prediction task tuned specifically for generated scFv sequences.

We demonstrate the use of deep learning approaches to infer thermostability attributes from scFv sequences generated experimentally and screened for their temperature sensitivity. First, with pre-trained language models (PTLMs), we assess the ability of unsupervised networks to predict thermostability with zero-shot and fine-tuned predictions. Then, we utilize supervised learning to train simple, predictive CNN architectures. To provide structural context to the supervised networks, we also feed the network with thermodynamic information via Rosetta energies. Further, we examine whether these networks could provide insights towards design of thermostable mutants, thereby improving biologics engineering. With this work, we present a proof-of-concept study of utilizing PTLMs and thermodynamic features towards relatively niche problems in protein informatics.

## Results

### Machine learning tasks can be tuned to identify scFv thermostability

The biological problem of thermostability prediction at the sequence-level involves identifying which sequences could result in a highly thermostable biomolecular structure; for antibodies and scFvs this implies conservation of the folded structural state and/or antigen binding upon high temperature stress (**Fig. 1A**). Thermal stability of proteins depends on residue-level biophysical attributes. But, for antibodies and scFvs, owing to the high consensus in sequences, deciphering heuristic or empirical rules based solely on sequence patterns for distinguishing thermostable and unstable sequences is challenging. Machine learning (ML) models have shown potential to extract higher-order relationships mapping sequences to function in the absence of underlying biophysical pathways, and they perform well on classification tasks. Leveraging sequence and structural information as features, ML approaches applied on a plethora of prediction tasks, for e.g., fluorescence landscapes (13–15), intrinsic stability (14, 16, 17), missense variant effects (18), protein fitness (19), antigen-specificity (20), have shown high success rates. With the availability of an explicit dataset with temperature-level information, we could extrapolate ML methods for thermostability prediction tasks.

**Fig. 1.**
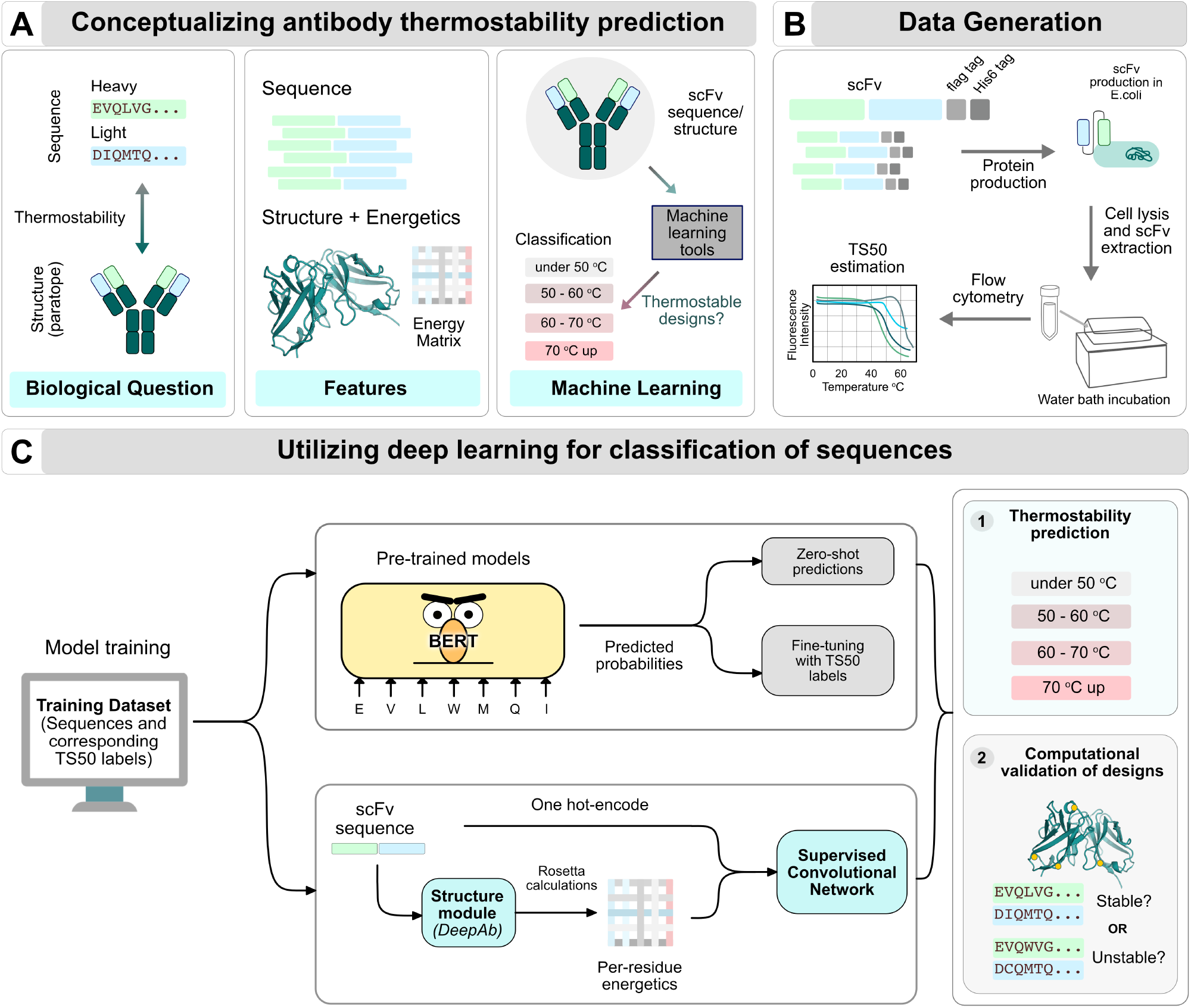
A pipeline to identify scFv thermostability using deep learning. (A) *The biological challenge of antibody thermostability prediction from sequences*. Antibody thermostability can determine the manufacturability of antibodies in downstream processes. The biological question that we aim to tackle involves given an scFv sequence, can one predict the thermal characteristics. Available data for this challenge can comprise of the amino acid sequences, structures, and calculated energetics. Leveraging antibodies with pre-determined temperature characteristics is paramount, however, the availability is scarce for such a dataset. (B) *Thermostability data generation*. To generate a dataset of scFv sequences with known temperature-specific features, we determined the loss of target binding of the scFv post high temperature stress to obtain a TS50 measurement. (C) *Training a classification network for predicting TS50 bins*. One of the approaches is transfer learning with unsupervised models (top branch). We utilized pre-trained BERT-like models (such as ESM1-b, ESM1-v, etc) to make (1) Zero-shot predictions and (2) Fine-tuned predictions with the labeled TS50 dataset. Another approach is to train a supervised model with calculated thermodynamic energies (bottom branch). We used sequence and structure-based features for supervised learning using simple convolutional models to train a classifier. The outcome of such trained ML models can be employed either for predicting thermostability of generated antibody sequences or to computationally validate experimental designs.

To learn temperature-specific contextual patterns in sequence-data, we sought to develop and train machine-learning models for thermostability prediction using scFv sequences. We collected temperature data from various engineering studies for developing thermostable single-chain variable fragment (scFv) molecules. The sequence data contained scFv sequences assembled by performing mutations to heavy and light chains from multiple germlines (Methods1 and Sup.Fig.S1). 2,700 scFv sequences from 17 germlines (further referred to as experimental sets) were collated to constitute the sequence data. Additionally, sequences from another scFv study (currently under clinical trials) and an isolated scFv dataset form out-of-distribution, blind test sets. For each sequence, thermostability is evaluated with a TS50 measurement representing the temperature at half-maxima of target binding, and this measurement serves as the temperature annotation (**Fig. 1B**). For the isolated scFv dataset, thermostability is evaluated with a Tm measurement representing the first transition from folding to unfolding as temperature is increased.

The temperature measurement (TS50) for 2,700 scFv sequences is used for two thermostability prediction tasks: (1) Regression: Prediction of TS50 measurement of a scFv sequence, and (2) Classification: Prediction of whether a given sequence corresponds to a thermally stable scFv. For the regression task, absolute values of TS50 measurements are used, whereas, for the classification task, the TS50 data are divided into four bins, namely under-50°C, 50°C-60°C, 60°C-70°C and 70°C-up. With these training data, we have trained two models (**Fig. 1C**). (1) Pre-trained language models (PTLMs), unsupervised BERT-like model architectures(21, 22), trained over large sequence corpus spanning evolutionary diversity. These models are trained to extrapolate learned representations of protein structures, function and biological activity at a sequence-level, and they can make zero-shot predictions or be fine-tuned with thermal stability data. (2) Supervised convolutional models, that equip simple deep-convolutional networks utilizing annotated thermostability data with feature encodings at a sequence-level (i.e. one-hot encoded amino-acid types), and an energy-level (i.e. thermodynamic energies obtained from putative three-dimensional structural models generated with DeepAb(23)). With both models, our aim is to predict thermostability of a given scFv sequence. By developing a model that can effectively filter and screen scFv sequences, we could significantly accelerate the identification of better variants for stable, manufacturable, multispecific biologics.

### Fine-tuning pre-trained sequence models with thermostability data improves classification performance over zero-shot predictions

Unsupervised models trained on a large corpus of protein sequences are reported to infer evolutionary relationships and statistical patterns about protein structure and function (22, 24). PTLMs have reportedly shown successful performance in downstream prediction tasks (e.g., predicting mutational landscapes, secondary structure and tertiary contacts(21)) without any additional supervision. To assess whether zero-shot learning from the PTLMs could be extrapolated for the thermostability tasks, we evaluate likelihood-based zero-shot predictions (22) from the ESM-1v (22) and UniRep (17) language models (**2A**). Next, we fine-tuned the pre-trained features from the ESM-1b and UniRep language models (Figure 2.B) specifically on our downstream thermostability prediction task to assess if that improves performance. **Figure 2C,E** show that zero-shot predictions do not in general correlate well with thermostability, either on TS50 data or on blind test sets. These results are contrary to those reported in prior work (22, 24, 25). It is important to note that the setting is quite different; prior work largely evaluates single mutations of a parent protein, whereas our datasets consist of multiple mutations (including insertions and deletions) and are derived from multiple parent proteins.

**Fig. 2.**
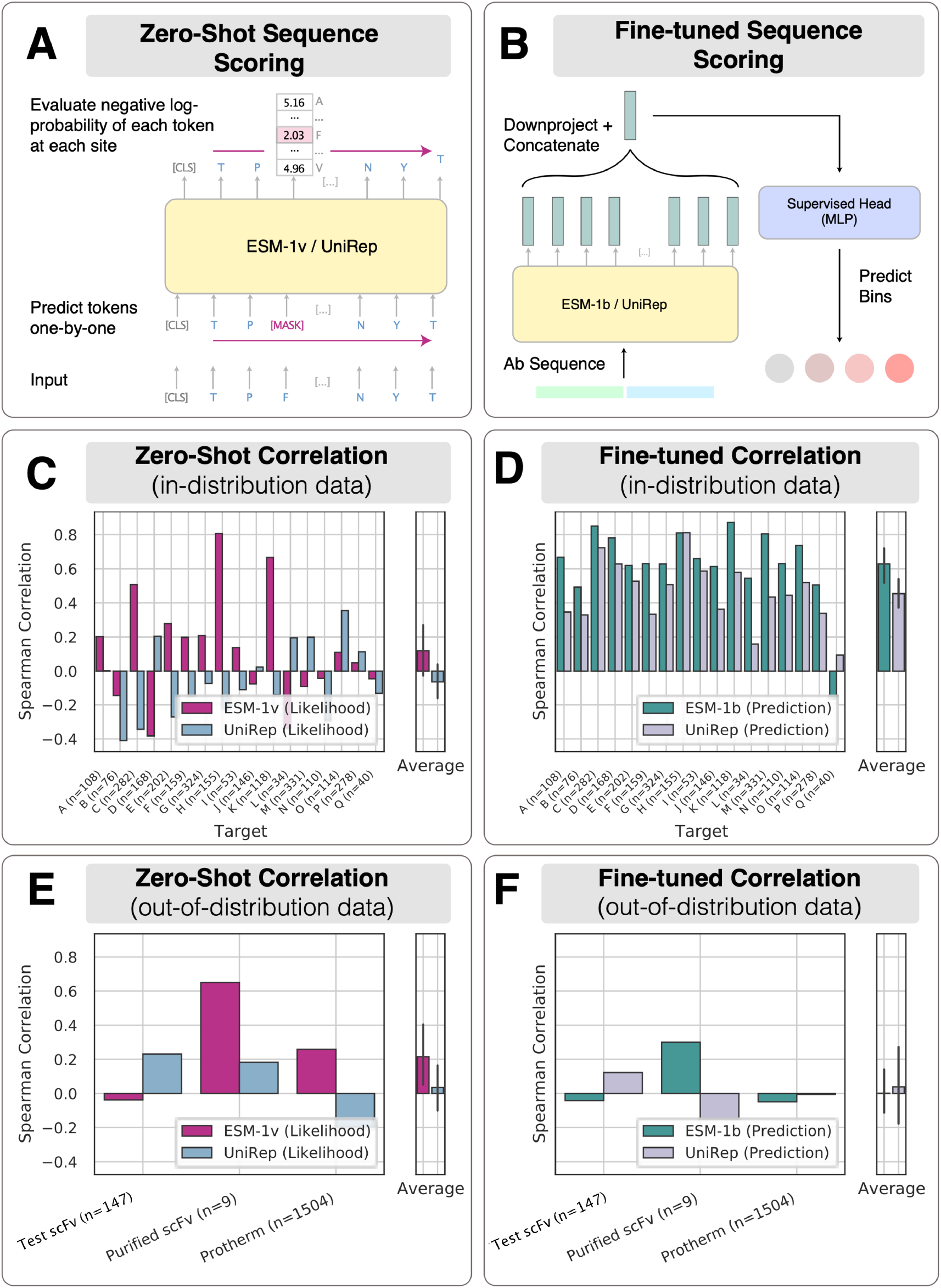
Fine-tuning over pre-trained unsupervised models improves correlation on withheld targets. (A) Zero-shot likelihood-based predictions with pre-trained models do not correlate strongly with the TS50 datasets. (B) Fine-tuning the pre-trained models on TS50 data from *n* − 1 targets significantly improves correlation on the heldout target. (C) Zero-shot likelihood-based predictions on blind test sets. (D) Models fine-tuned on TS50 data do not generalize well to blind test sets.

Fine-tuned predictions from both ESM-1b and UniRep achieve moderate to high average Spearman correlation on held-out targets when trained on TS50 data (0.63 and 0.45 respectively) (**Figure 2D**). However, these predictions do not generalize well to blind test sets (**Figure 2F**). This suggests there is some underlying structure in the sequences in the TS50 set that the model can exploit to make predictions, but which does not generalize to new datasets.

### Supervised network trained with energy features improves generalizability across out-of-distribution datasets

Unlike pre-trained models utilizing large sequential information, Shanehsazzadeh *et al*. demonstrated that small supervised models could achieve competitive performance on downstream prediction tasks benchmarked in TAPE (15). Similar predictive performance was also reported for antigen-specificity prediction with supervised convolutional networks (20). We built a supervised convolutional model with the scFv sequences for the thermostability prediction task. **Figure 3A** shows the detailed architecture of our supervised CNN deep-learning model. Since the sample size of the experimental dataset was relatively small (2,700 scFv sequences), we decided to supplement the network with structure-specific information. To feed the network a structural context, we incorporated the energetics as a two-dimensional *i*-*j* residue energy matrix. For each scFv sequence, we generated a structural model with DeepAb (23) and evaluated the thermodynamic features (total energy split into one-body, *i*-*i*, and two-body, *i*-*j*, residue energies) using Rosetta ref2015 (26) energy function. The contributions of *i*^*th*^ residue with every *j*^*th*^ residue (where *j* ∈ 1, *N* such that *N* = total number of residues) were tabulated and binned in an *i*-*j* matrix that constituted the energy features. Finally, we equipped the model with two branches : (1) Sequence branch with one-hot encoded amino-acid sequences. (2) Energetics branch with *i*-*j* residue energy matrix. The final model architecture and hyper-parameters are reported in **Figure 3A**, and this model was trained and evaluated with the available sequences (Sets A-Q). The model architecture was built such that contributions from either of the two branches could be turned off to obtain sequence or energy dependence over the classification performance.

**Fig. 3.**
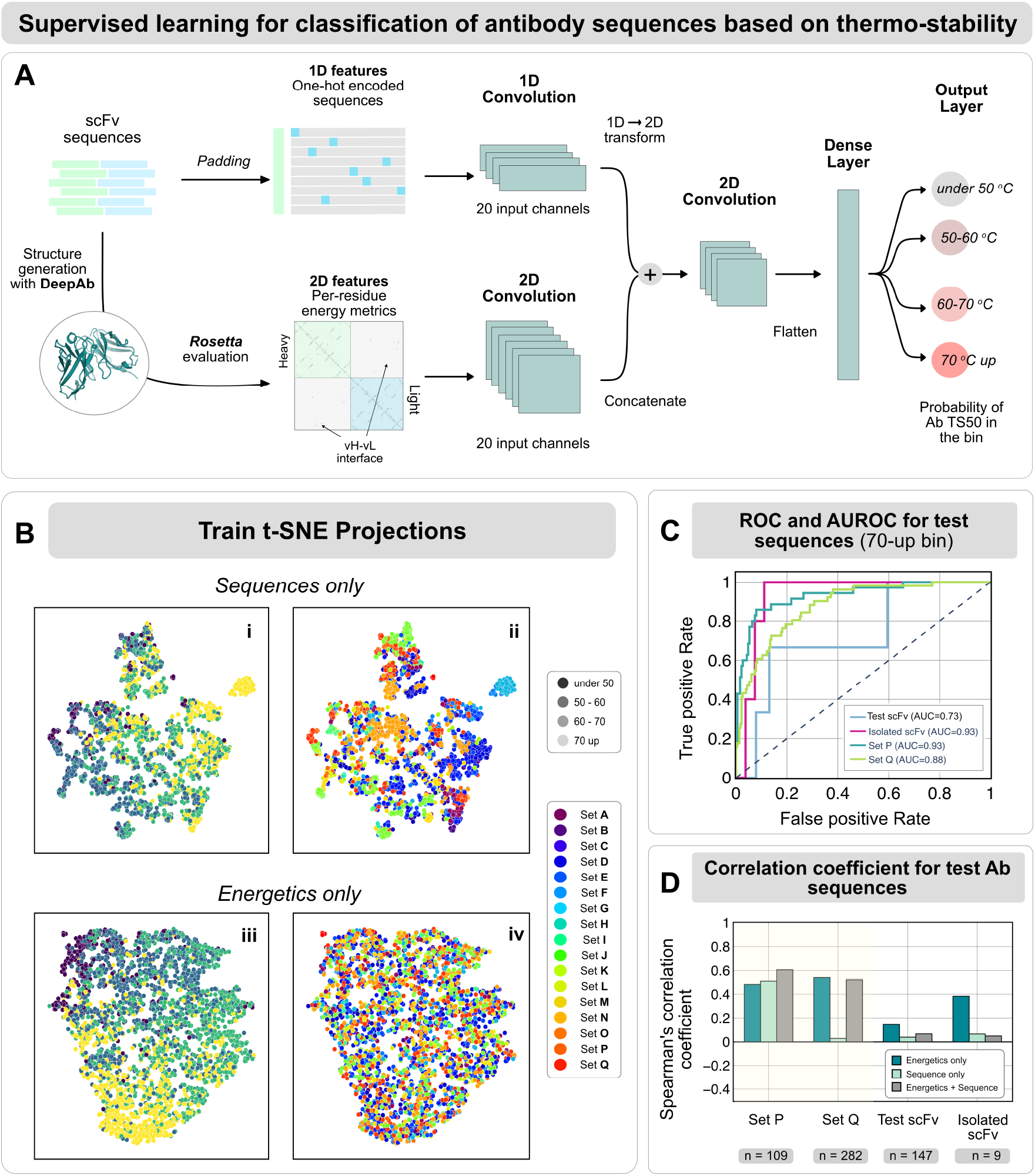
Energy features can extract ‘generalizable’ information of thermostability. (A) The supervised convolutional network architecture for classification of antibody sequences. The input scFv sequences pass a structure-generation module with DeepAb followed by Rosetta-based evaluation to estimate per-residue energies for each amino acid residue in the scFv structure. The sequences are one-hot encoded (top branch) and the energetic features, represented as an i-j matrix(bottom branch), are provided to the network. The output from the sequence branch and the energy branch are concatenated and flatten to pass through a dense-layer and the probabilities of the sequence to lie in each of the temperature bins is output. (B) t-stochastic neighbor embeddings from the energetics-only model colored by the temperature bins. (C) Receiver-operating characteristic curve to demonstrate the classification of the test sequences for the above-70 bin with the energetics-only model. Note that Test scFv and Isolated scFv have a smaller sample size, explaining the relatively less rugged nature of the curves. (D) The model’s performance metrics for the classification task on completely blind test scFv sequences is reported with the Matthew’s correlation coefficient for four test cases

In spite of the sequence diversity in the experimental data, we wanted to investigate whether there was an underlying relationship between the sequences; whether the experimental sets from which the sequences were derived had an impact over prediction accuracies. We analyzed the representation learned by the sequence-only model and the energetics-only model by projecting the embeddings from the dense layer for each sequence into two dimensions via t-distributed stochastic neighbor embedding (t-SNE) **Figure 3B**. The sequence-only model embeddings were clustered by their experimental set, as evident by the aggregation of colored points in **Figure 3B.iv**. On other hand, the energetics-only model embeddings were independent of any clustering based on the experimental set as demonstrated by the noisy embedding for energetics (**Figure 3B.ii**). Thus, in spite of a sequentially diverse dataset, fine-tuned and supervised models trained only on sequence-features are able to infer the underlying experimental origin of the sequences and skew thermostability predictions, making them less generalizable towards newer, blind datasets.

Since the energetics-only model was relatively more generalizable, we assessed the performance of this supervised model by constructing a receiver-operating-characteristic (ROC) curve derived from the prediction of the 70-up bin (**Figure 3C**). As we aim towards identifying thermostable sequences, the prediction accuracy of the 70-up bin is most important. We evaluated the ROC for four test datasets: two held-out (Sets P and Q) and two blind datasets representing a test scFv and an isolated scFv. The area under ROC is over 0.7, denoting a high classification accuracy. **Figure 3D** shows the Spearman’s correlation coefficient for all four test datasets, with the energetic-only, sequence-only and energetics+sequences models respectively. On held-out datasets (Set P and Q), the coefficients are over 0.5 for energetics-only model with energetics+sequences model showing an equally better performance. But on blind datasets, the performance drops for energetics+sequences and sequences-only (coefficients under 0.1). The energetics-only model still shows relatively higher correlation for the blind datasets (0.2 and 0.4 respectively).

Finally, as a control, we randomly initialized the weights in the SCNN for the classification task and found that it is unable to distinguish sequences based on thermostability (**Sup.Fig.S1**). Further, on the test sets, we performed weighted random predictions i.e. we predicted the classification bin label with a weighted random choice, with sample size in each bin set as the weights (**Sup.Fig.S2-S3**). In both these tests, the energetics-only SCNN were able to decipher some relationship between the energetics of the scFv and the thermostability. The randomly initialized models could not demonstrate any discernible relationship demonstrating the significance of learned representations from supervised data.

### Networks trained with experimental TS50 data can distinguish thermal stability-enhancing designs

With improved thermal stability predictions on scFv sequences, we aspire to optimize and engineer antibody or scFv molecules for specific biomolecular applications. Thus, we sought to evaluate the ability of our predictive models in discriminating between thermostable and thermally degenerate mutations. Prior studies by Koenig *et al*.(27) and Warszawski *et al*.(9) detail thermal aggregation experiments on point mutations for an anti-VEGF antibody (PDB ID: 2FJG/2FJF (28)). In both the studies, a deep mutational scanning (DMS) experiment was performed for the antibody, and selected point mutations that improved binding enrichment over wildtype were analyzed for their fragment antigen-binding (Fab) melting temperature (T_*m*_). These point mutants (20 mutations compiled from both the studies) serve as a test case to evaluate whether the networks trained on TS50 temperature measurements could obtain insights about related temperature-dependent attributes such as thermal aggregation; and whether they distinguish the thermally-enhancing and thermally-hampering mutations. Although the model is better suited to classify diverse sequences rather than point mutations, with this test, we wanted to understand the generalizability of our models trained on scFv sequences and TS50 measurements to antibody sequences and melting temperature measurements.

To perform an unbiased analysis, we performed point mutations over the antibody (a computational DMS), and analyzed the classification performance of our PTLM models (ESM-1v, fine-tuned) and our SCNN model (ensemble of energetics-only CNNs) on these point mutations. **Fig. 4** compares our 70-up bin predictions with the experimental thermostable mutants for the heavy and light chains with the two models. The 20 mutation positions that were validated experimentally are highlighted as spheres in the cartoon representations. Our networks identify five out of the 20 mutations correctly (highlighted in magenta). Surprisingly, all five of these mutations comprise the framework residues. Further, for 18 out of 20 mutations, the SCNNs could identify the residue position correctly, albeit predicting different amino-acid mutations as most thermostable. Out of 4540 point mutations analyzed (*N*_*res*_ = 227 residues, 20 amino acids per residue), experimental data was available for only 20 point mutations. Since only 0.44% of the total possible mutations in the anti-VEGF antibody were assessed for melting temperatures experimentally, the validation dataset for thermostability is scarce. Further, in spite of being temperature-specific attributes, TS50 and T_*m*_ are different experimental measurements and do not correlate exactly. It is, therefore, remarkable that our networks could predict the thermostable residue positions in 90% of the cases, with 25% successful predictions (correct residue positions as well as amino-acid residues). By extrapolating the networks trained on TS50 measurements over alternative thermal aggregation experiments (T_*m*_ in this case), we demonstrate that intrinsic thermal attributes could be captured by such models. Moreover, on comparing the residue positions violating the germline consensus sequence for the anti-VEGF Ab, we observed different amino acid mutations, highlighting the ability of these models to provide mutations orthogonal to traditional germlining approaches (**Sup.Fig.S5-S6)**. With a more diverse and larger training dataset, it would be possible to develop a more robust model. Our results suggest that these networks could serve as a useful tool for screening or filtering scFv (or even antibody) sequences for temperature-specific antibody design pipelines.

**Fig. 4.**
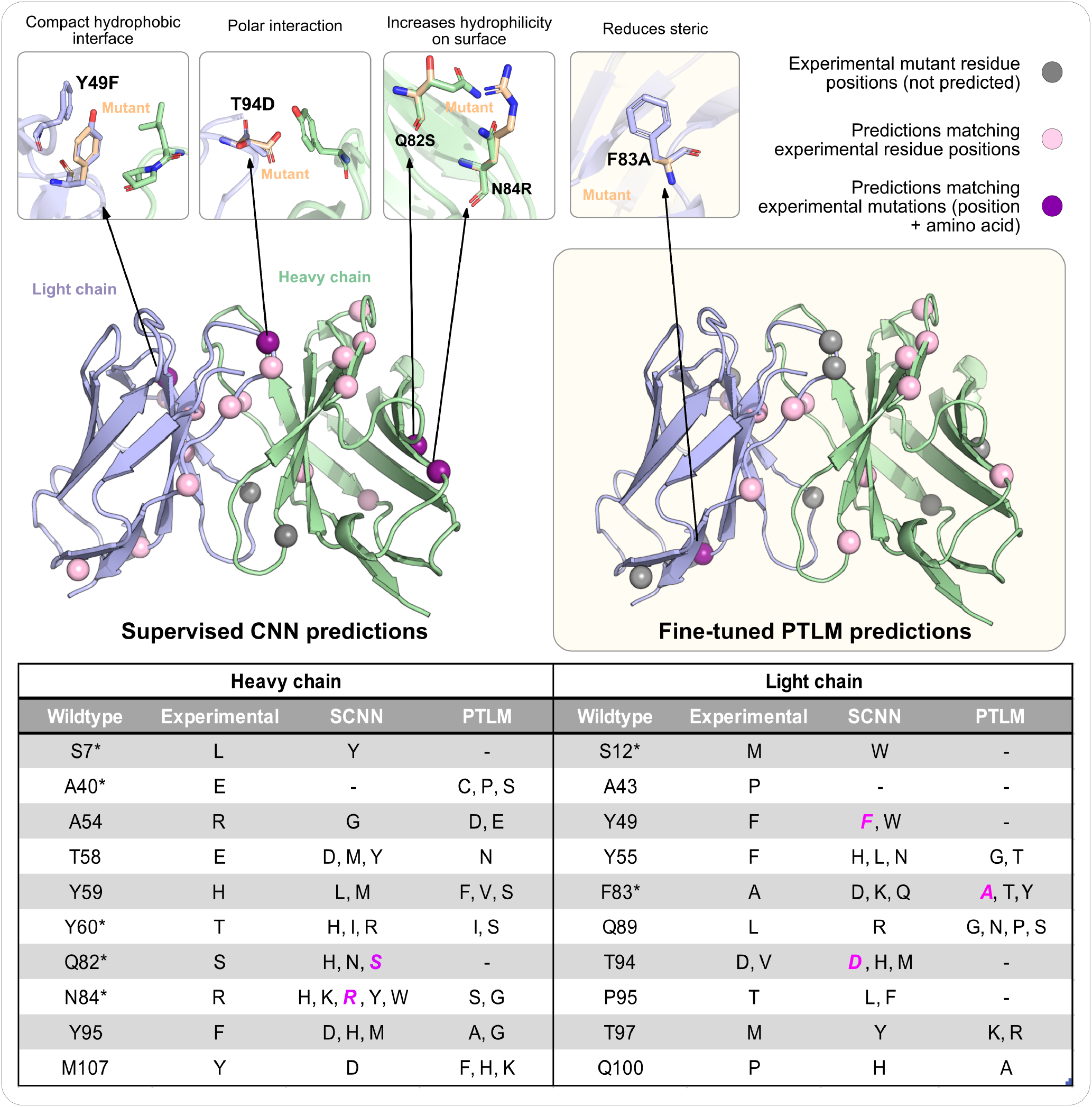
Computational deep mutational scan of an antibody variable fragment shows agreement with experimental thermal denaturation data. Validating all the point mutants with our supervised network for anti-VEGF antibody (PDB ID: 2FJG(bound) and PDB ID: 2FJF (unbound)), we observed synergies in mutants predicted in the over-70ºC bin and the experimental thermal denaturation data available from prior work (cite Warsawski2019 and Koenig2017). Spheres indicate the experimentally validated mutants that improved Tm; pink indicates predictions from the network with the same residue position, but different amino acid mutation; red indicates the predictions matching experimental data and gray indicates mutations which were not observed in computational predictions. Thumbnails highlight the mutations in agreement with experiments and potential interactions. The table illustrates the comparison with the experimental and computational predictions.

## Discussion

Thermostability is an important determinant of developability. To address the limitations in developing thermostable biological candidates, antibody engineering efforts are directed towards identifying and screening for sequences that can improve thermostability. In this work, we have tested two approaches for prediction of thermostable scFv sequences from features learned with a sequential and thermodynamic context. As the corpus of sequence databases is vast (billions of sequences from diverse protein families), we equipped the unsupervised learned representations via pre-trained language models to classify sequences into temperature-specific bins quantifying their thermostability. Unlike conventional machine-learning approaches that use sequence or structural-coordinate features, we incorporated enriched information with thermodynamic features. Further, we tested the performance of using energetic features on small, supervised CNN models for the classification tasks. Finally, we demonstrated the applicability of our work for antibody engineering efforts by identifying experimentally validated melting temperature (Tm) enhancing mutations on an anti-VEGF antibody. While the primary objective of this work was to study proof-of-principle for scFv thermostability classification with machine-learning models, the secondary objective was to identify ‘generalizable’ feature representations that can aid in creating a pipeline for rapid, computational screening and validation of scFv and antibody sequences based on their thermal characteristics.

First, we extrapolated the zero-shot learning and fine-tuning principles of large-scale PTLMs for inferring temperature-dependent biophysical attributes of scFv sequences. We acknowledge the limitations of extrapolating language models trained on massive sets of protein sequences to scFv-specific sequence data. Unlike natural proteins which evolve under selective or evolutionary pressures, antibodies are often selected for binding towards a particular antigen. Hence, models trained on a huge corpus of natural proteins might be unsuitable for capturing scFv or antibody sequence information. We anticipate that by employing an antibody-specific language model (for eg. sequences from the Observed Antibody Space (29)) as the pre-trained network, we could better explore the biophysical attributes for scFv and antibodies.

To learn generalizable representations, we sought to utilize enriched structural context of scFv sequences with thermodynamic (energy) features. We focused on the residue-wise energy features for each putative structure generated from the scFv sequence, and used these energy-dependent features to train our CNN model. Validation of this small, supervised CNN network demonstrates the ability of energy-features to be more generalizable and predictive towards thermostability. For the relatively small dataset of 2,700 scFv sequences, we found that thermodynamic context could infer biophysical attributes better. Note that our training data is non-uniform, with some experimental sets skewed largely towards higher temperature bins owing to the selection procedure for generating the scFv sequences. With stringent selection and screening of sequences for uniform distribution in the temperature bins, along with incorporation of negative data (i.e. variant sequences that did not express, showed drastically low TS50 values, or unfolded at room temperature), we could generate well-distributed datasets for training our ML models. Additionally, utilizing accurate three-dimensional antibody structures could refine the energetic input and potentially improve performance. Better quality of experimental and structural data would thus provide a better insight towards predicting thermostability.

In spite of the networks being trained for classification tasks, there are potential avenues in extending these models towards biologics engineering and design. With this work, we demonstrated how we could use this classification network to filter and suggest thermostability-enhancing point mutations. Although the temperature point mutant dataset was fairly scarce, our models demonstrated substantial predictive accuracy towards mutants with higher thermal aggregation (Tm). One might argue that our network is trained on TS50 measurements, and so evaluating mutants with improved melting temperature is not plausible. However, as both the metrics evaluate the thermal attributes of the sequence, we can extend the patterns learned by our networks over alternative temperature-dependent data (i.e. thermal aggregation). With this work, even with training on sparse experimental data, we want to highlight the use of ML models towards evaluating an essential biophysical characteristic. Energy features represent a refined, information-rich resource that can add thermodynamic context which ML models are often deprived of. In closing, we have demonstrated a proof-of-concept of using PTLMs and SCNN architectures for thermostability prediction. Moreover, these models can also be equipped with experimental information derived from alternate physical properties (e.g. viscosity, binding enrichment), thereby enabling the engineering and design of antibodies and broad-scale biologics.

## Methods

### Experimental Methods

#### Generation of scFvs

scFvs with a (G4S)3 linker were cloned as a single construct into a pTT vector with a puromycin selection marker. Constructs were transfected into a mammalian CHO-K1 cell line and stably expressed at a 4 mL scale. After 21 days post transfection, VCD and viability were measured and the expression level of secreted proteins in conditioned medium were analyzed by non-reduced SDS PAGE gel. Cells were further incubated with magnetic beads coupled with either proA (for scFvs with lambda variable domains) or proL (for scFvs with kappa variable domains) overnight. The beads were separated from cell media and following by washed with PBS for three times and water for 2 times. scFvs were eluted from the magnetic beads with a low pH buffer (100mM glycine, pH2.7) and neutralized with 3M Tris (pH11). Differential Scanning Flourimetry (DSF) was carried out to determine the melting temperature of the purified material. Briefly, molecules were heated at 1.0°C/min on a nanoDSF instrument. Changes in tryptophan fluorescence were monitored to evaluate protein unfolding and aggregation. The Tm is reported as the midpoint between the unfolding onset and the max unfolded state.

#### TS50 Screening Assay

The thermostability of scFvs was screened by determining the loss of target binding after high temperature stress. To this end, soluble scFvs (VH-(G4S)3-VL) containing a C-terminal FLAG-tag (DYKDDDDK) and a 6xHis-tag were produced in *E*.*coli* TG1 (Agilent, Santa Clara, USA) in 10 mL LB cultures. Protein production was induced with 1 mM IPTG. Bacteria were then centrifuged, and the cell pellet was resuspended in 1 mL Gibco™ DPBS. Cells were lysed with 4 freeze/thaw cycles and residual cells and cell debris were removed by two centrifugation steps. 100 *μ*L of these crude extracts were transferred into 0.2 mL tubes and subjected for 5 min to different temperatures in waterbaths (4°C, 50°C, 60°C, 70°C). After incubation, the tubes were directly transferred on ice and human target transfected CHO-cellswere incubated with 50 *μ*L of the lysates. Bound scFvs were detected and analyzed by flow cytometry. Median fluorescence intensity values were determined and plotted. The temperature corresponding to half maximal binding of each scFv was calculated (TS50). The scFv sequences were further binned into sets based on the identity of the antigen they bind (not random). Note that since the sets were not curated for a ML task, there is lack of an uniform distribution across sets.

#### nanoDSF T_m_ Method

Thermal melting (T_*m*_) temperatures were determined by running a Trp Shift Study on the Prometheus, NT.48. A thermal ramp was applied at 1.0°C/min with start temperature 25°C and stop temperature with 95°C. Unfolding was measured by the fluorescence ratio 350nm/330nm. Data analysis and T_*m*_ determination was performed using PR. ThermControl v2.0.4. Samples were normalized to 1.0 mg/mL in formulation buffer prior to Tm analysis.

### Pretrained language models

We evaluate three pretrained language models on their ability to predict stability of scFv sequences. The first model, UniRep (17), is an mLSTM (30) with 1900 hidden units pretrained on the Pfam database (31). Following the “evotuning” methodology proposed by the authors, we collect MSAs for each sequence in our TS50 set, combine all sequences into a single dataset, and further pretrain the model on this evolutionarily related set of sequences using the implementation from (32).

Additionally, we consider both the ESM-1b (21) and ESM-1v (22) transformer models. Both are 33 layer, 650M parameter transformer models, pretrained with masked language modeling on the Uniref database (33). The primary difference is that ESM-1b is trained on a 50% sequence identity filtered dataset (Uniref50), while ESM-1v is trained on a 90% sequence identity filtered dataset (Uniref90). ESM-1v is specifically designed to improve zero-shot likelihood evaluation of protein sequences.

#### Zero-shot evaluation

One approach to predicting stability with pretrained language models is to directly use model likelihood or pseudolikelihood (22, 24). Sequences which are more likely under a model are predicted to be more stable. Suppose *x* = *x*_1_*x*_2_ … *x*_*n*_ is a protein sequence with each *x*_*i*_ representing a residue. UniRep models the probability of each residue given all preceding residues. As a result the likelihood of a sequence can be efficiently evaluated as

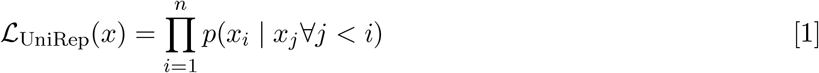

ESM-1v models the probability of masked residues given unmasked residues. It is not possible to efficiently decompose this probability and obtain an exact likelihood. However, it is possible to obtain the pseudo-likelihood of a sequence:

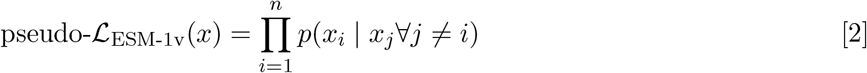

In practice, the log-likelihood and pseudo-log-likelihood are evaluated for numerical stability. Additionally, ESM-1v comes as an ensemble of five models trained with different random seeds. The predictions from all five models are averaged to obtain the final pseudo-log-likelihood.

#### Finetuned evaluation

The other approach to predicting stability with pretrained language models is to finetune a task-specific model using supervised data. For the UniRep model we the methodology suggested by the authors and take the final hidden state along with the average of previous hidden states as a fixed-length vector representation of 3900 hidden units. For the ESM-1b model, we follow methodology suggested by (34) and downproject each per-residue representation to 4 dimensions, followed by a concatenation. This results in a fixed length embedding of size 4*L*, where *L* is the maximum sequence length in the TS50 dataset. If a sequence has length less than *L*, it is padded with zeros.

These embeddings are passed through a linear layer with a hidden dimension of 512, followed by tanh activation, then to a final layer to predict class logits. Parameters of the UniRep and ESM-1b models are frozen during training. Parameters of the head model (including the initial downprojection for ESM-1b) are trained with the Adam optimizer and a learning rate of 10−3.

For TS50 data, models are trained on all but one target and evaluations are made on the held out target. For non-TS50 data, an ensemble of TS50 models (one for each holdout target) is used to make predictions.

## Supervised models

### Dataset Curation

#### Sequence inputs

Datasets for TS50 measurements of scFvs from all experimental sets were aggregated to form a single dataset. The scFv sequences comprised of a heavy and a light chain linked together with a Glycine-Serine (G4S)_*x*_ linker. We chose to create a dataset with the scFv sequences, split into their respective heavy and light chain sequences, and instead of classifying sequences based on their thermostability, we included their TS50 measurements. The distribution of the scFv sequences across the experimental sets and the test dataset is illustrated in Sup.Fig. S1. Sets P and Q were removed, along with the sequences of the test scFv and the isolated scFv to constitute the held-out set. The amino acid sequences were one-hot encoded to form an input of dimension, (*V*_*H*_ + *V*_*L*_ + 3) × 21, where *V*_*H*_ and *V*_*L*_ correspond to the heavy and light chain sequences respectively. The additional token to the amino acids’ one-hot encoding corresponds to the delimiter at the start and end positions of the scFv sequence, and between heavy and light chains to indicate a chain-break.

#### Energetic inputs

To obtain the energetics input, the sequences were first passed through a structural module .i.e. the DeepAb protocol for antibody structure prediction. For each predicted structure, we ran a Rosetta Relax and refinement protocol for side-chain repacking (XML scripts in the Supplementary). Energy estimation in Rosetta starts with a energy relaxation step to reduce steric clashes (Rosetta Relax) with constraints to the start coordinates so that the accuracy of backbone structure (predicted by DeepAb) is not diminished. The all-atom model is refined further with 4 cycles of side-chain packing to obtain a robust structure and the lowest energy structure is chosen for further calculations. For each refined model, we then estimated the residue-residue interaction energies with the residue_energy_breakdown application. We converted these one-body and two-body energies to a two-dimensional *i*-*j* matrix that served as the energetic information for training in the supervised CNN models. The energy values in the *i*-*j* matrix were also further binned together in 20 bins between the lower-end and upper-end energies of [−25, 10] REU respectively. We included an additional bin for the start, end and chain-break tokens respectively. The dimension of the pairwise energy data is this, *L* × *L* × 21.

### Model Architecture

We evaluated supervised convolutional networks with sequence, energetics and sequence+energetic features to predict thermostability of scFv sequences. We use the scFv sequences to predict the structure (DeepAb (23)) and obtain energetic features with Rosetta. All the sequence and energy input is converted to a fixed length embedding of size *L* and *L* × *L* respectively, where *L* represents the maximum sequence length in the dataset, such that *V*_*H*_ and *V*_*L*_ are the maximum lengths of the heavy and light chains respectively. The sequences less than *L* are padded with zeros. While padding the sequence and energy embedding are fed to two parallel branches of the model, one with a 1D convolutional layer and other with a 2D convolutional layer. The sequence and energetic input is fed to the network, such that sequences pass through a 1D CNN and energies pass through a 2D CNN, followed by concatenation (. The output is then passed through another 2D convolutional layer, and then a final layer to get the logits. We perform a softmax over the logits to obtain the class probabilities. The parameters of the supervised model are trained with Adam optimizer with categorical cross entropy (CCE) loss and a learning rate of 10^−3^.

To estimate the predicted TS50 value (.i.e. the regression task), the probabilities are weighed with the mean TS50 value in each bin. For the prediction of the temperature bins for a given sequence (prediction task), an *argmax* over the probabilities gives the expected thermostability (temperature bin).

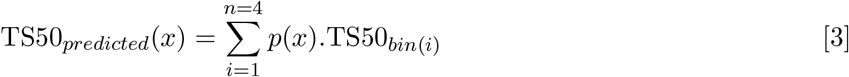

Alternatively, we performed additional tests with different CNN architectures for the energy branch (1D-CNN by flattening the *i*-*j* matrix and 2D-CNN with absolute energy values, i.e., no binning). The architectures for these additional models was optimized by performing a randomized search for the parameters and variables, layers, dropout, batch size, number of filters, kernel size, strides, epochs and pooling size. The performance was assessed with Spearman’s correlation coefficient and we found out that the 2D-CNN binned architecture for the energy branch worked better than other architectures. The 1D-CNNs resulted in loss of individual interactions as the average contribution by most of the residues is similar, it was difficult for the network to discern an useful context. For 2D-CNNs with absolute data, as better sequences are identified with lower, negative energies, the information was lost in the convolutions. 2D-CNN binned architecture resolves both the issues; it provided context of individual residue-residue interactions and the binning ensured that relevant energy information is retained through the convolutional layers. A comparison of these methods is further illustrated in the Sup.Fig.S2. As the CNN models are trained on a smaller dataset, to reduce variance we ensemble three CNN models trained on different sets to obtain an ensemble of CNNs, which we use for our predictions with the anti-VEGF antibody thermal denaturation data.

### Extrapolating trained predictive models for design

The anti-VEGF DMS dataset was generated to enrich binding by designing of multi-point variants. To determine whether our predictive models have potential in protein design, we created a computational DMS on the anti-VEGF antibody (PDB ID: 2FJG). Each residue position in the sequence was mutated to 19 other amino acids to obtain mutant sequences. Each sequence was one-hot encoded to obtain the sequence data, and the energetics dataset was generated following the procedure mentioned prior. This sequence and energy input was fed to the models and the point mutants classified in the 70-up temperature bin by our predictions were cross-verified with the experimental results.

## Data Availability

The source code for the PTLM and supervised CNN networks, along with the Rosetta scripts, will be made available prior to publication. The experimental training data and the trained weights are classified and property of Amgen Inc.

## ACKNOWLEDGMENTS

This work was supported by XXXX.

## References

1. C Spiess, Q Zhai PJ Carter, Alternative molecular formats and therapeutic applications for bispecific antibodies. Mol. immunology 67, 95–106 (2015).

2. X Zhong, AM D’Antona, Recent Advances in the Molecular Design and Applications of Multispecific Biotherapeutics. Antibodies (Basel, Switzerland) 10 (2021).

3. M Klinger, J Benjamin, R Kischel, S Stienen, G Zugmaier, Harnessing T cells to fight cancer with BiTE® antibody constructs–past developments and future directions. Immunol. reviews 270, 193–208 (2016).

4. J Dong, et al., A stable IgG-like bispecific antibody targeting the epidermal growth factor receptor and the type I insulin-like growth factor receptor demonstrates superior anti-tumor activity. mAbs 3, 273–288 (2011).

5. GL Moore, et al., A robust heterodimeric Fc platform engineered for efficient development of bispecific antibodies of multiple formats. Methods (San Diego, Calif.) 154, 38–50 (2019).

6. MS Sawant, CN Streu, L Wu, PM Tessier, Toward Drug-Like Multispecific Antibodies by Design. Int. J. Mol. Sci. 21 (2020).

7. BR Miller, et al., Stability engineering of scFvs for the development of bispecific and multivalent antibodies. Protein engineering, design & selection : PEDS 23, 549–557 (2010).

8. A Goldenzweig, et al., Automated Structure-and Sequence-Based Design of Proteins for High Bacterial Expression and Stability. Mol. cell 63, 337–346 (2016).

9. S Warszawski, et al., Optimizing antibody affinity and stability by the automated design of the variable light-heavy chain interfaces. PLoS Comput. Biol. 15, 1–24 (2019).

10. MDS Kumar, et al., ProTherm and ProNIT: thermodynamic databases for proteins and protein-nucleic acid interactions. Nucleic acids research 34, D204–6 (2006).

11. L Jia, R Yarlagadda CC Reed, Structure based thermostability prediction models for protein single point mutations with machine learning tools. PLoS ONE 10, 1–19 (2015).

12. Y Yang, et al., Pon-tstab: Protein variant stability predictor. importance of training data quality. Int. J. Mol. Sci. 19 (2018).

13. KS Sarkisyan, et al., Local fitness landscape of the green fluorescent protein. Nature 533, 397–401 (2016).

14. R Rao, et al., Evaluating Protein Transfer Learning with TAPE. Adv. neural information processing systems 32, 9689–9701 (2019).

15. A Shanehsazzadeh, D Belanger, D Dohan, Is Transfer Learning Necessary for Protein Landscape Prediction?, 1–10 (2020).

16. GJ Rocklin, et al., Global analysis of protein folding using massively parallel design, synthesis, and testing. Science 357, 168–175 (2017).

17. EC Alley, G Khimulya, S Biswas, M AlQuraishi, GM Church, Unified rational protein engineering with sequence-based deep representation learning. Nat. methods 16, 1315–1322 (2019).

18. VE Gray, RJ Hause, J Luebeck, J Shendure, DM Fowler, Quantitative Missense Variant Effect Prediction Using Large-Scale Mutagenesis Data. Cell Syst. 6, 116–124.e3 (2018).

19. C Hsu, H Nisonoff, C Fannjiang, J Listgarten, Learning protein fitness models from evolutionary and assay-labeled data. Nat. Biotechnol. (2022).

20. DM Mason, et al., Optimization of therapeutic antibodies by predicting antigen specificity from antibody sequence via deep learning. Nat. Biomed. Eng. 5, 600–612 (2021).

21. A Rives, et al., Biological structure and function emerge from scaling unsupervised learning to 250 million protein sequences. Proc. Natl. Acad. Sci. 118 (2021). 439

22. J Meier, et al., Language models enable zero-shot prediction of the effects of mutations on protein function. Adv. neural information processing systems 35 (2021).

23. JA Ruffolo, J Sulam, JJ Gray, Antibody structure prediction using interpretable deep learning. bioRxiv, 2021.05.27.445982 (2021).

24. TA Hopf, et al., Mutation effects predicted from sequence co-variation. Nat. biotechnology 35, 128–135 (2017).

25. AJ Riesselman, JB Ingraham, DS Marks, Deep generative models of genetic variation capture the effects of mutations. Nat. methods 15, 816–822 (2018).

26. RF Alford, et al., The Rosetta All-Atom Energy Function for Macromolecular Modeling and Design. J. Chem. Theory Comput. 13, 3031–3048 (2017).

27. P Koenig, et al., Mutational landscape of antibody variable domains reveals a switch modulating the interdomain conformational dynamics and antigen binding. Proc. Natl. Acad. Sci. United States Am. 114, E486–E495 (2017).

28. G Fuh, et al., Structure-function studies of two synthetic anti-vascular endothelial growth factor Fabs and comparison with the Avastin Fab. The J. biological chemistry 281, 6625–6631 (2006).

29. A Kovaltsuk, et al., Observed Antibody Space: A Resource for Data Mining Next-Generation Sequencing of Antibody Repertoires. The J. Immunol. 201, 2502–2509 (2018).

30. B Krause, I Murray, S Renals, L Liang, Multiplicative lstm for sequence modelling in 5th International Conference on Learning Representations. pp. 2872–2880 (2017).

31. A Bateman, et al., The pfam protein families database. Nucleic acids research 32, D138–D141 (2004).

32. EJ Ma, A Kummer, Reimplementing unirep in jax. bioRxiv (2020).

33. BE Suzek, H Huang, P McGarvey, R Mazumder, CH Wu, Uniref: comprehensive and non-redundant uniprot reference clusters. Bioinformatics 23, 1282–1288 (2007).

34. NS Detlefsen, S Hauberg, W Boomsma, What is a meaningful representation of protein sequences? arXiv preprint 2012.02679 (2020).

