## Supplementary Information for "Towards generalizable prediction of antibody thermostability using machine learning on sequence and structure features"

[a]Department of Chemical and Biomolecular Engineering, The Johns Hopkins University, Baltimore, MD 21218, USA.; [b]Electrical Engineering and Computer Science, University of California, Berkeley; [c]Therapeutic Discovery, Amgen Research (Munich) GmbH, 8 Munich, Germany; [d]Department of Therapeutics Discovery, Amgen Research, Amgen Inc., Thousand Oaks, CA 94080, USA; [e]Department of Therapeutics Discovery, Amgen Research, Amgen Inc., South San Francisco, CA 94080, USA

Figs. S1 to S7

### 1. Supplementary Results

#### A. Nature of the dataset.

We compiled scFv sequences and TS50 measurements from 17 different experimental sets, such that each set represent a germline sequence mutated to obtain single-point or multi-point variants. Each of the scFv sequences were screened at high temperature stress and their TS50 measurements are recorded. We want to highlight that the experimental dataset was non-uniform and potentially skewed towards the higher temperature bins (i.e. 60-70 and 70-up bins). A distribution of the training, validation and test datasets is illustrated in **Sup.Fig.S1**. The sequence data representation is such that the green regions highlight high consensus and red regions indicate poor consensus. We can observe that the $Gly_4$/Ser linker region between the $V_H$ and $V_L$ chains is evident. For our training, the heavy and light chain sequences were separated from the linker. Sup.Fig.S1.B highlights the temperature distribution of the TS50 measurements to show the skewed nature of the experimental dataset.

#### B. Ensemble models result in robust predictions.

For supervised CNN models, we showed that the energy models can learn embeddings that separate the proteins based on their thermostability characteristics as opposed to the sequence models which are skewed towards the experimental sets. Before selecting a 2D-CNN binned model for energetics however, we performed an assessments of alternative ways to feed the energy data, namely, a 1D flattened input and a 2D-input with absolute residue-wise energy values. We tested these architectures to validate if any of these methods perform better than sequence only model, and result in a better separation in embeddings. The results are illustrated in **Sup.Fig.S2**. For the energetics only case, 2D-CNN with binned inputs perform better than the other two cases and so this architecture for chosen for training.

To reduce the variance in the model performance, we chose to ensemble performance of three models by averaging their predictions and generating an ensemble of CNNs. With the same architecture, we trained the supervised CNNs on different dataset splits. We shuffled the experimental sets (experimental sets A to Q) to obtain 3 different pairs of training and held-out data. **Sup.Fig.S2.B,C** demonstrate the slightly improved performance with the Ensemble of CNNs. In order to test a control, we employed a randomly initialized CNN and compared the embeddings generated with this randomized model with the ensemble of CNNs. Clearly, our model can better separate the sequences based on their thermostability characteristics, even with limited sequence size and training diversity. We believe that utilizing thermodynamic energies as features for machine learning objectives has further potential for alternative bio-physical characteristics prediction tasks.

We also test our performance on blind datasets, i.e. the test scFv and the isolated scFv sequences, with respect to a weighted random prediction model. For this case, we bias the randomization with the weights of the sample size of each bin as observed from the training dataset. This sample size is then used as a probability with the random number generator to predict the bins of the sequences. We acknowledge that the data points are limited, non-uniform and heavily skewed, yet we were able to observe some trends as opposed to the weighted random predictions. **Sup.Fig.S3** shows a confusion matrix highlighted with the probabilities of the prediction in each bin. Here, we want to highlight that even though our model could not predict the exact bins of the sequences, we could observe some trends. The predictions in the top-most bin, are skewed more towards the higher temperature regions in our predictions as opposed to the weighted random predictions. This implies that, while predicting blind sequences, if the top-most bin i.e. 70-up bin, is considered, then there is a higher probability of actually selecting sequences which are thermostable i.e. lie in the top 2 bins, 60-70 or 70-up. This is important as it can help us remove redundant, potentially less thermostable sequences and with a well-curated training sets, simple supervised networks could be useful for making robust design estimations.

##### C. Performance on individual experimental datasets.

Finally, we show an overall head-to-head performance for all the test sets with the PTLMs and Supervised networks on in-distribution (training and validation) and out-of-distribution (test) datasets. Supervised networks have a higher Spearman's correlation coefficient for all the training and validation sets, as well as the two blind sets. The average correlation coefficient on validation sets is 0.6 and on test sets is 0.4 which shows acceptable classification characteristics. With this study, we want to highlight the potential of using PTLMs and Supervised CNN networks for thermostability prediction task and lay the groundwork for utilzing energetics as feature encodings in future deep learning work.

#### 2. Supplementary Methods

##### A. Supervised CNN Model architecture.

A detailed schematic of the supervised convolutional networks is illustrated in **Sup.Fig.S7**. As described prior, we supply the network with one-dimensional sequence input and two-dimension pairwise energetic input. Both the sequence and energy branches pass through a single convolutional layer with Batch Normalization and ReLU activation. Next, the sequence input is transformed and concatenated with the energetic input. For training sequence-only or energetics-only model, the other branch is switched off and a tensor of zeros is passed instead. For sequence + energetics model, the concatenation is performed as illustrated. The concatenated matrix is then passed through another 2D convolutional layer, flattened and supplied to a dense layer to output logits for each class. The class probabilities are further obtained by performing a softmax over the logits.

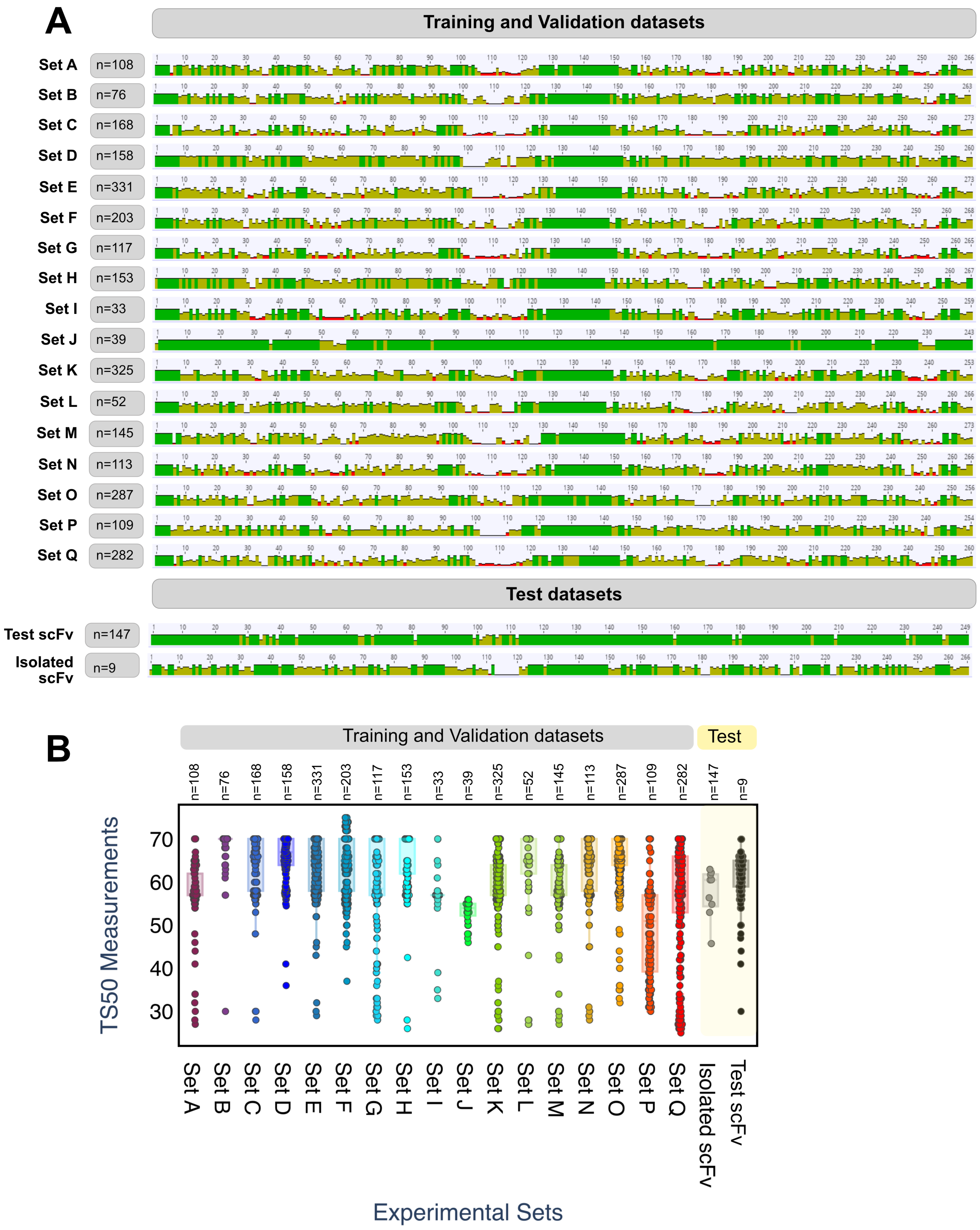


**Fig. S1. Data distribution of the TS50 dataset.** (A) Plots of percent identity of aligned dataset sequences per position for each of the training, validation, and test datasets. Green and full height bars indicates 100% identity in the aligned residue position while red and low height bars indicate low percent identity in the aligned residue position. (B) Distribution of the experimental TS50 values for each of the training, validation, and test datasets. For the isolated scFv test set, experimental $T_m$ values were measured and plotted here on the same axis as TS50 for comparison.

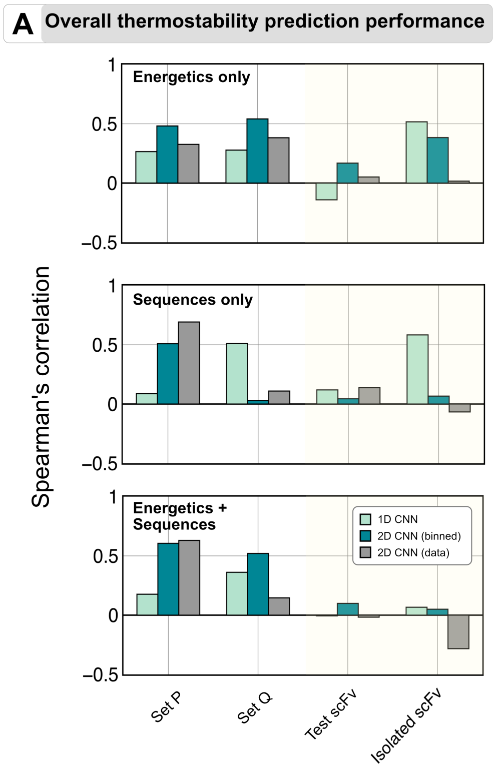


**B** Spearman's ρ for models (Energetics only)

| Models | Validation | Set P | Set Q | Test-Ab | Isolated scFv |
|---|---|---|---|---|---|
| 1D CNN | 0.81 | 0.28 | 0.27 | -0.142 | 0.52 |
| 2D CNN (binned) | 0.77 | 0.54 | 0.48 | 0.17 | 0.38 |
| 2D CNN (data) | 0.19 | 0.38 | 0.33 | 0.051 | 0.17 |
| Ensemble | 0.86 | | | 0.15 | |

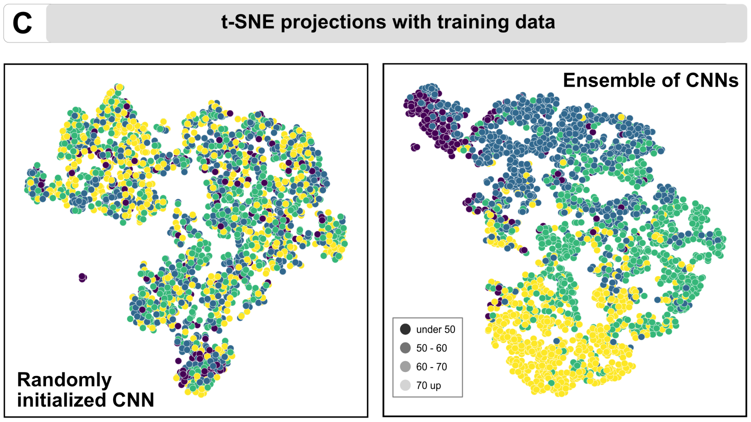


**Fig. S2. Comparison of all features for the supervised models.** (A) Overall thermostability prediction performance for energetics-only, sequences-only and energetics+sequences. (B) Comparison of spearmann rho for different architectures of the energetics only model. (C) t-SNE projections for a randomly initialized CNN and an Ensemble of CNN model.

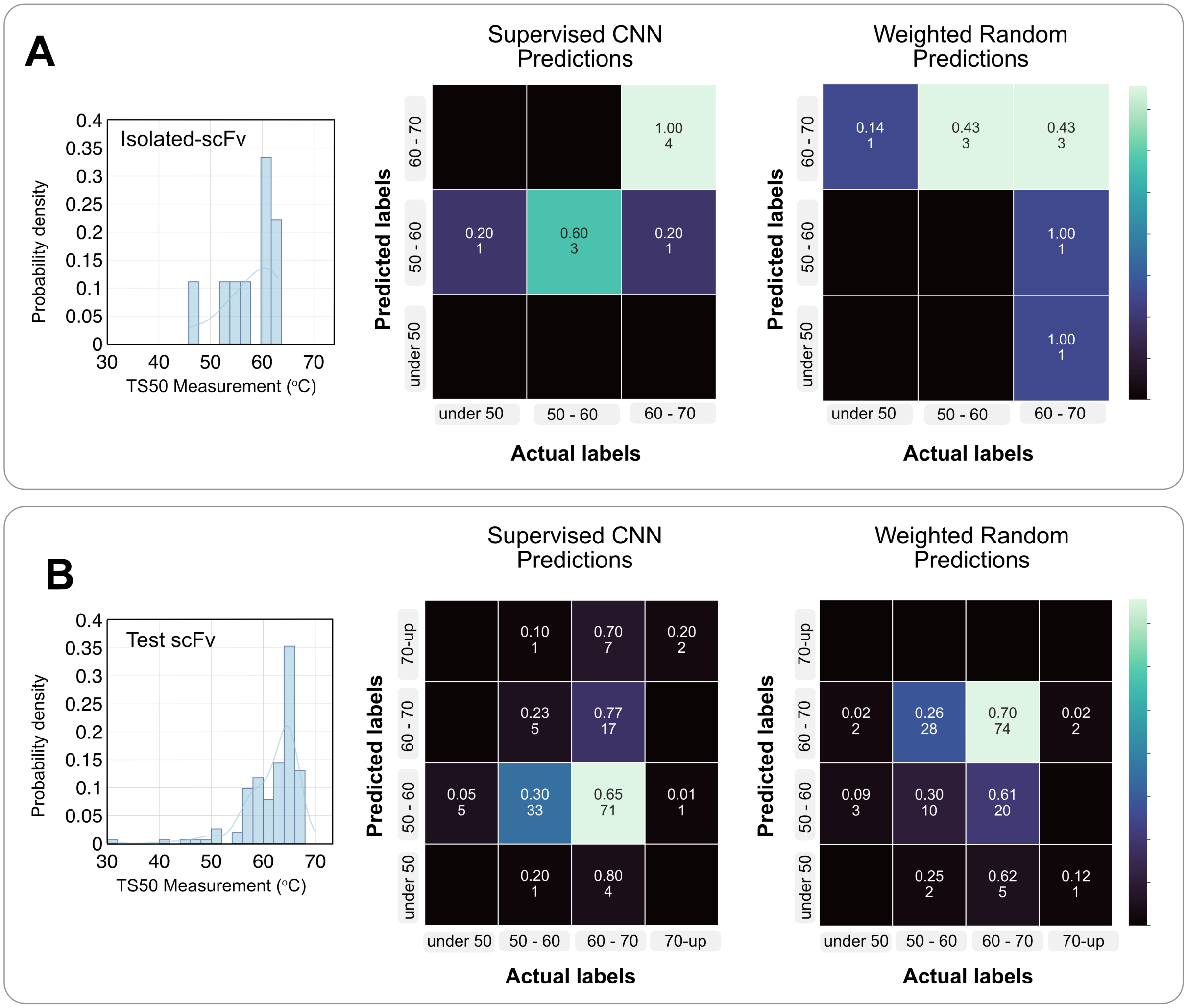


**Fig. S3. Distribution of TS50 performance with scFv sequences to show the range of sequence length that is incorporated and to explain poor performance for the test scFv case.** Performance of (A) Isolated-scFv and (B) Test-Ab, with SCNN and a weighted-random prediction model.

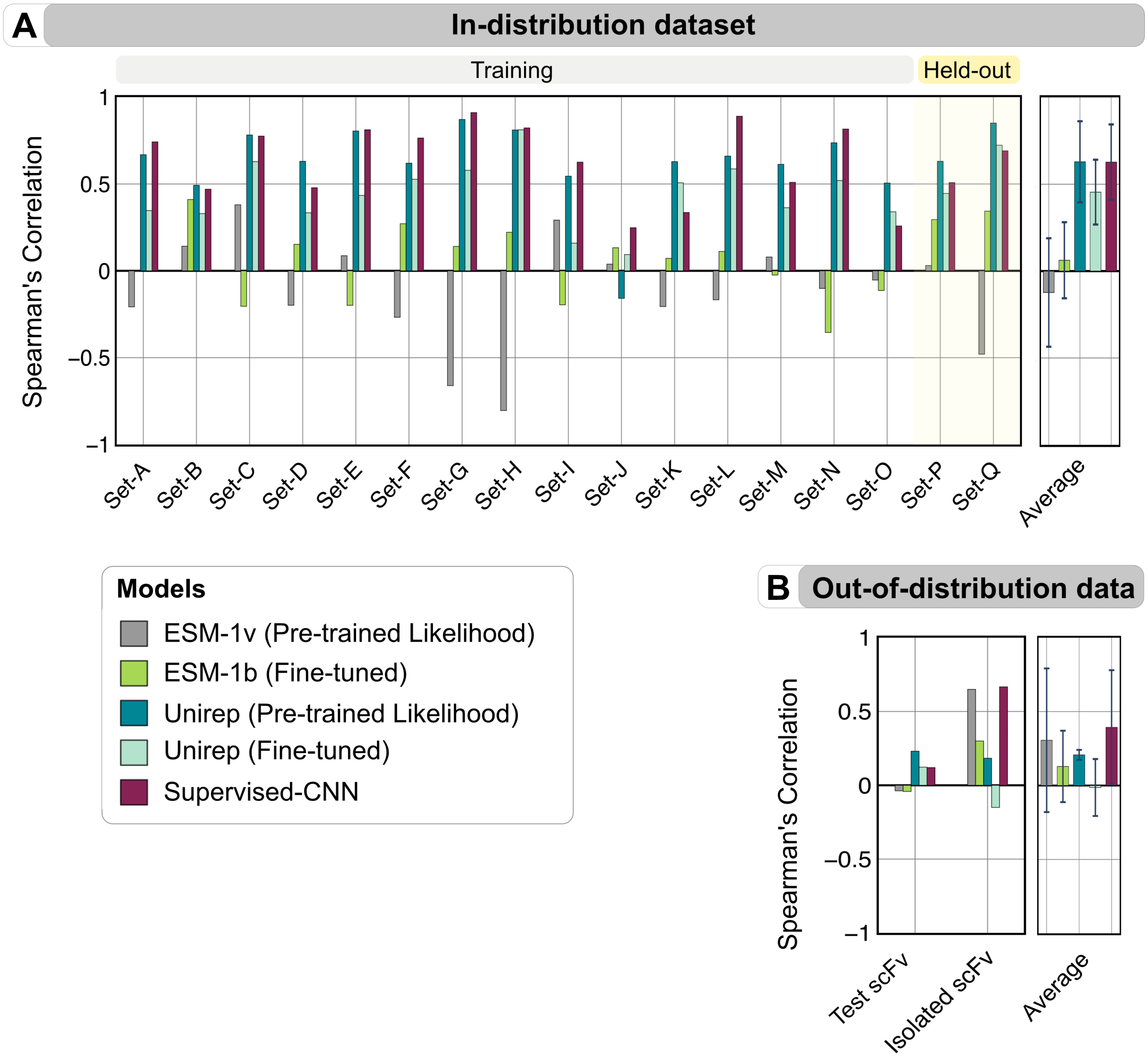


**Fig. S4.** **Overall performance of all models on the training and testing dataset.**

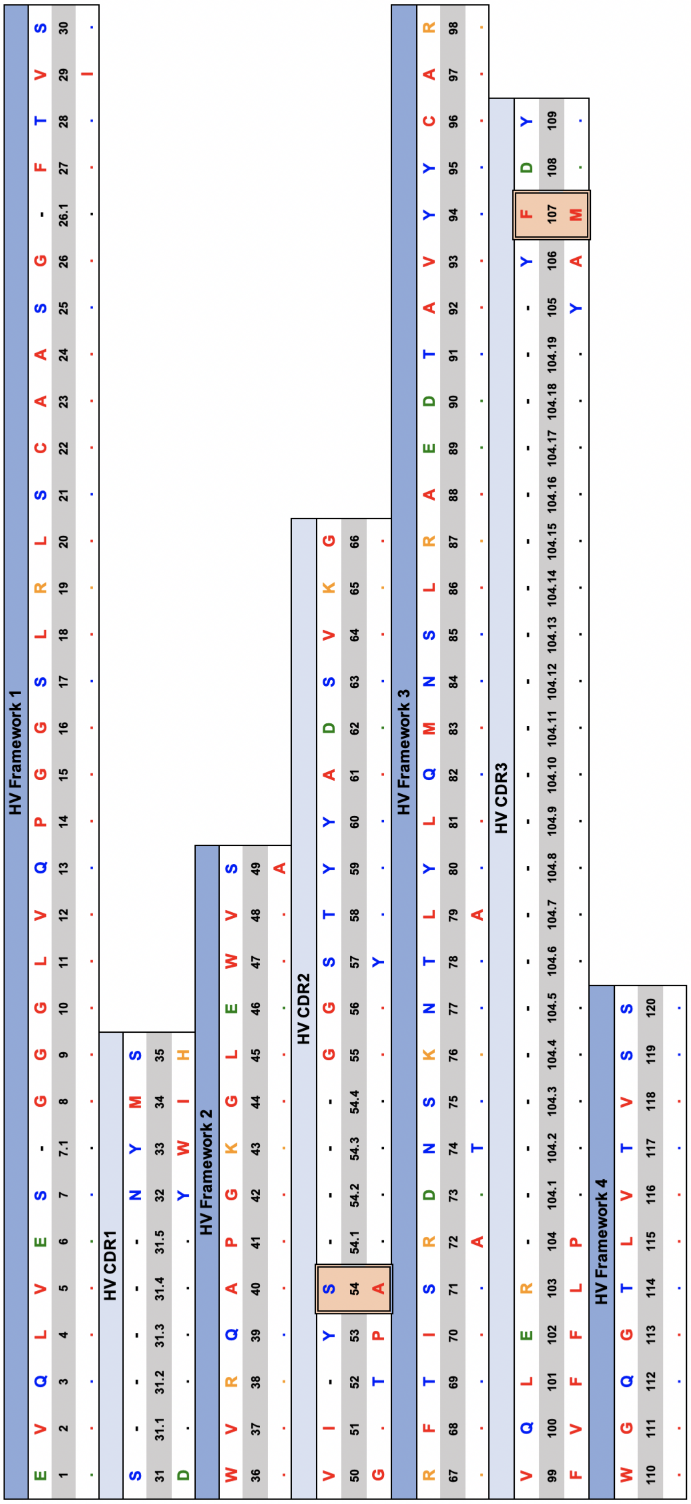


**Fig. S5. Alignment of the G6-antibody (PDB ID: 2FJG) with the germline for the heavy chain.** (top) Germline sequence (bottom) Amino acid sequences in the antibody. Numbers denotes the linear residue numbering for the Ab. Highlighted box represents the residue position/s that were identified by prior studies. (*Koenig et. al*, 2017 and *Warszawski et. al*, 2019)

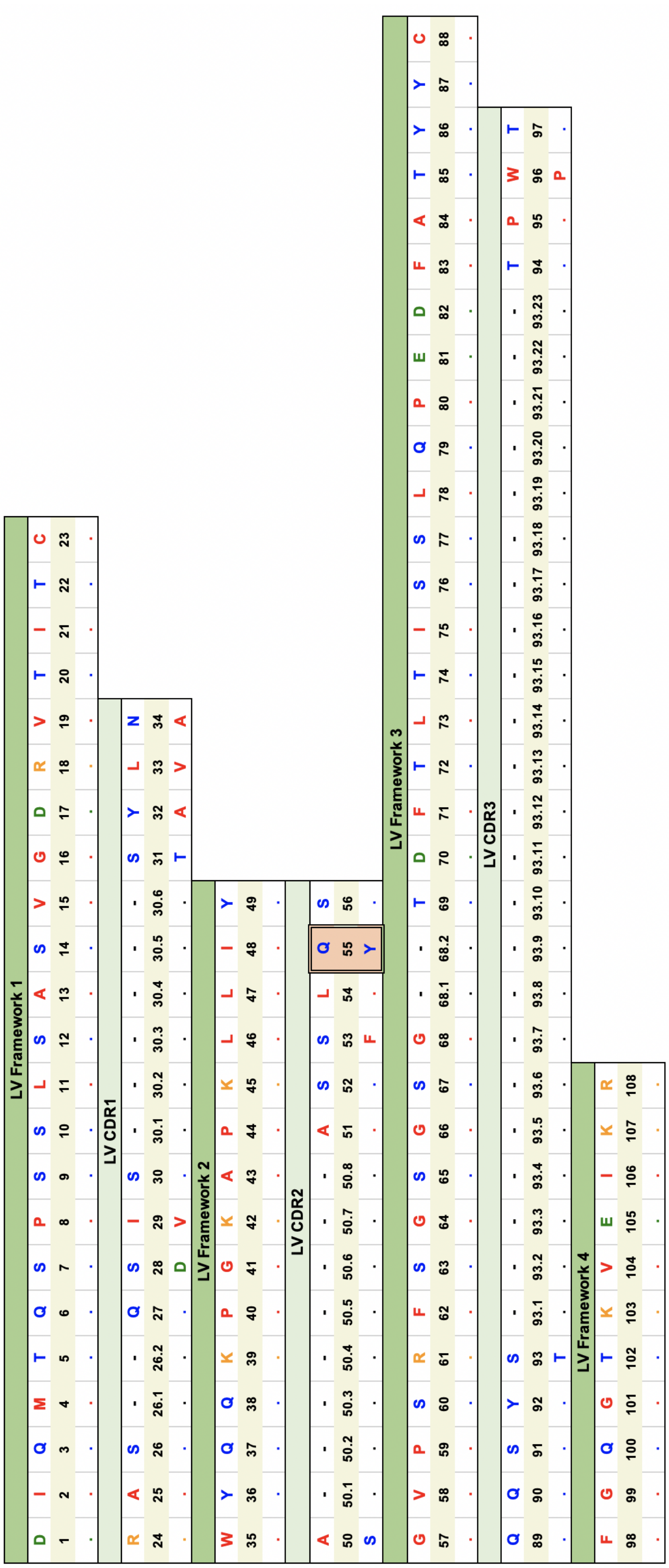


**Fig. S6. Alignment of the G6-antibody (PDB ID: 2FJG) with the germline for the light chain.** (top) Germline sequence (bottom) Amino acid sequences in the antibody. Numbers denotes the linear residue numbering for the Ab. Highlighted box represents the residue position/s that were identified by prior studies. (*Koenig et. al*, 2017 and *Warszawski et. al*, 2019)

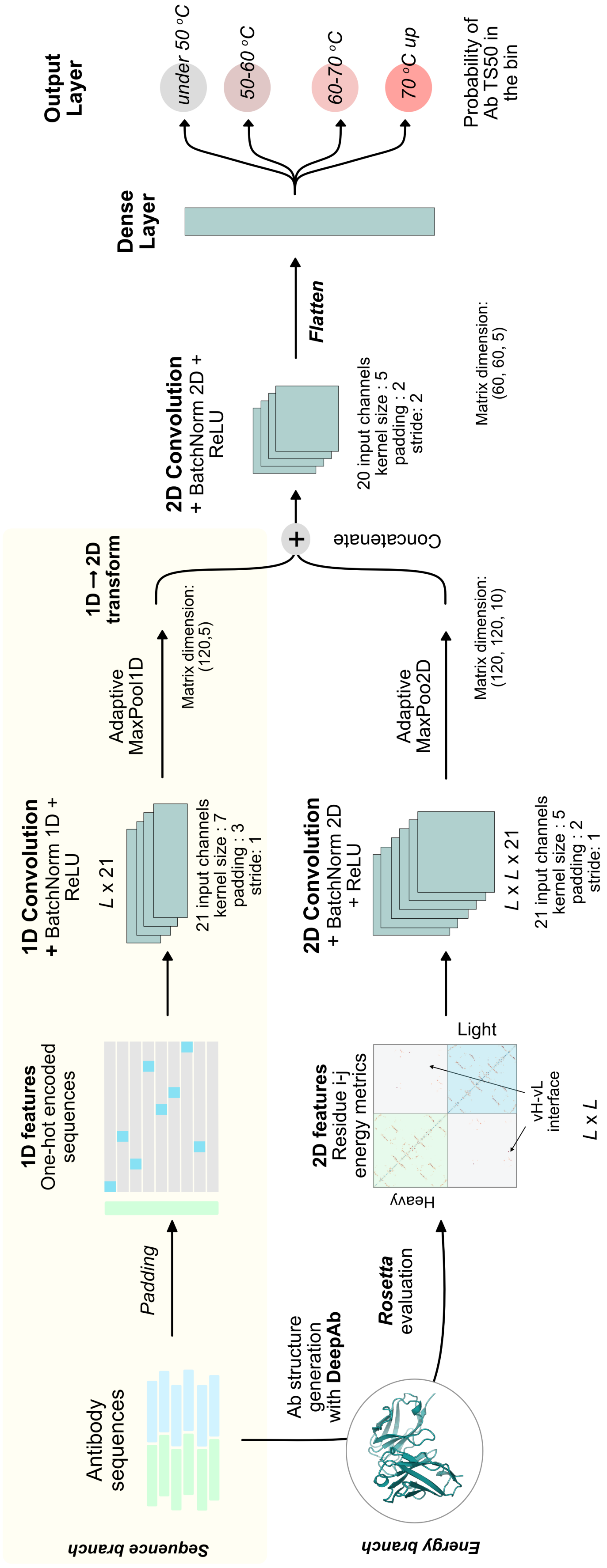


**Fig. S7. Schematic of the Supervised CNN with parameters and layers.**
